# Protein adlayer thickness on colloidal nanoparticle determined by Rayleigh-Gans-Debye approximation

**DOI:** 10.1101/852228

**Authors:** L Yuan, Z Zhai, L Chen, X Ge, D Li, G Ge

## Abstract

Reference materials (RM)-assisted Rayleigh-Gans-Debye approximation (rm-RGDA) has been developed and used to *in situ* determine the size and thickness of the adlayer on the particles in solution. The particle size determined by rm-RGDA is quite close to that measured by electron microscopy but significantly smaller than that measured by DLS. The BSA adlayer absorbed on PS50, PS100 and SiO_2_ NPs is 3.3, 0.9 and 1.2 nm, respectively, and close to those observed by SEM, which is 4.6, 1.3 and 3.8 nm, respectively. The FTIR analysis results show that the BSA absorbed on larger particles or hydroxyl-abundant surface, e.g. PS100 and SiO_2_ NPs can lose its secondary structure, e.g. *α*-helix, to a great extent and that absorbed on a more curve surface, e.g. smaller PS50 particles can largely preserve its secondary structure as its free state. The measurement results show the curvature of the NPs is closely related to the structure change of the adsorbed protein. This method provide a facile and new approach to measure the size and its adlayer change of the hybrid and core-shell structured nanoparticles in a wide range of wavelength.

**SIGNIFICANCE:** Quantitative study on the adsorption of the protein on colloidal nanoparticles is an important approach to understand the biophysical effect, compared with other ex situ methods such as TEM and SEM, where the specimen are undergone pre-processing and no longer the original state in measurement. It is, therefore, a big challenge. In order to cope with this challenge, UV-vis based RGDA has been developed and applied to in situ measure the size of the dispersed colloidal nanoparticles and their protein adlayer thickness, where the protein adlayer thickness on the colloidal nanoparticles can be easily determined. We believe this method provide a facile and sensitive way to in situ measure the dimension change of hybrid colloidal nanoparticles.

## INTRODUCTION

Nanoparticles and their adlayer can be characterized by many technologies such as light scattering, electron microscopy and differential sedimentation centrifuge etc.(1–8) The most obvious and simplest method for analysing shape, size, and size distribution of submicron particles is electron microscopy. However, the method is costly, laborious, micro area detection and skill-demanding and it involves the analysis of particles outside their actual dispersion medium and is often subject to sample perturbation and preparation artefacts. Moreover, it requires taking and inspecting many electron micrographs to gain a representative result. Other kinds of high resolution microscopy, such as the scanning tunnelling microscopy or the atomic force microscopy, or the transmission electron microscope which can resolve structures at nanometer or sub-nanometer scale, are even more expensive and complex to deal with. Currently, dynamic light scattering (DLS) is the most popular method for measuring size of submicroscopic particles. However, the DLS instrument can only irradiate and detect the scattering signal at a fixed wavelength and specific angle, which limits its use to measure the colloidal particles in much diluted state and those that can absorb the photons at this wavelength. Moreover, in the presence of excess ligand, e.g. protein, the size and its distribution determined by DLS significantly deviates from the actual ones. As an alternative, UV-vis spectroscopy based spectral method is considered as a less costly, more accessible, and potentially faster method to study the size feature for the non-absorbing colloidal particles. Complimentary to DLS, UV-visible spectrum-based scattering method can measure the colloidal nanoparticles in a direction of 180° along the irradiation angle, where the signal that can be detected by UV-vis scattering technology is much stronger than DLS. In addition, the choice for the irradiation wavelength to optimize its signal-to-noise is quite broad compared with DLS. The UV-vis scattering method provides a powerful and non-destructive tool for real time detection of the thermal degradation of PLLA due to its very sensitive to minute colour changes for the PLLA melt.(9) Khlebtsov et al have demonstrated the practical application of this simple method to estimate the average size of liposomes and their thickness.(10) Leslie Greengard et al have extended the similar method to electromagnetic scattering to the time-domain.(11) The experimental results demonstrate that if the scattering spectrum of a suspension and the weight concentration of dry substances comprising liposome particles, by the gravimetric method, are known, one can estimate the thickness of the particle shell.

The theoretical base of this method is Rayleigh-Gans-Debye approximation (RGDA) and the relevant discussion will be seen in the following section. Elsayed et al have used Rayleigh-Gans-Debye approximation (RGDA) to analyse the experimental turbidity spectra of extruded unilamellar vesicle suspensions to derive vesicle size and obtain comparable result like dynamic light scattering.(12) Guschin and Glasse have determined the nanoparticle size distribution by similar spectroscopic method and, consequently, to characterize the material properties.(13, 14) Cournil et al have demonstrated the potential and limits of the turbidimetry based on UV-vis absorbance as a particle sizing method in both theoretical and experimental aspects.(2) Shmakov and Blevennec et al have studied the factors that influence the quantitative measurements and limit of the classical turbidity spectrum and light scattering method for nanoparticle solution.(3, 15) However, this method is less successful to in situ measure the size and adlayer thickness of the protein absorbed colloidal particles directly without purification and pre-treatment of the sample.

In order to cope with this problem, a modified RGDA model has been developed and optimized, to measure the size of the nanoparticles and the thickness of the protein adlayer on the particles, respectively, assisted by the use of certified reference materials (CRM), e.g. 80 nm of PS nanoparticles. The measurement of the protein adlayer on the NPs by DLS is usually disturbed by the presence of the free protein molecules in solution and, therefore, over-estimates the size of the particles and thickness of the adlayer, while the modified RGDA method can treat this sample quite well and give rise to more accurate and reliable results like TEM/SEM.

## MATERIALS AND METHODS

### Theoretical Model

Rayleigh scattering works well for the particles in nanoscale when 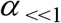 and the particle size is smaller than 1/10 wavelength (***α*** = ***πd/λ***, where *d* and *λ* is the characteristic length (diameter) and wavelength of the light, respectively).

For a particle scatterer, the irradiation intensity (*i_ϕ_*) in any scattering direction can be expressed as Eq. 1 according Rayleigh-Gans-Debye approximation.

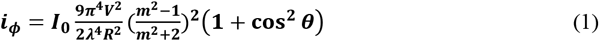

where *V* is the particle volume, *R* is the distance from the scatter, *λ* is the irradiation wavelength, *m* is the relative refractive index of the particle to the solvent (see the Supporting Information) and *I_0_* is the incident irradiation intensity. The protein refractive index (RI) used here can be estimated by Schuck’s method through the determination of its RI increment, *dn/dc* for unmodified protein based on their amino acid composition (see the Supporting Information) and the RI for PS, SiO_2_ and water is calculated according to responding empirical formula(see the Supporting Information). The total scattering intensity in all direction can be integrated around the scattering sphere with the radius of *R* as Eq. 2

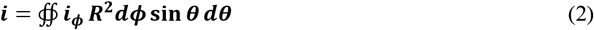

If the suspension contains *N* particles and the shape of the particle is sphere, the total scattering intensity (*I_S,t_*) can be expressed as Eq. 3.

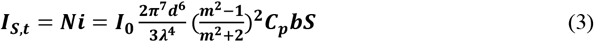

where 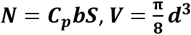, and *b*, *S, C_p_* are the optical path length, illumination spot area and number concentration of the scatterers, respectively.

### Measurement Model

The scattering intensity (***I_S_***) of the particles in solution is determined by a conventional UV-Vis spectrometer and can not be measured straightforward like absorbance (***I_A_***). When the sample is irradiated by a light beam with an incident intensity of ***I*_0_**, it can be expressed as

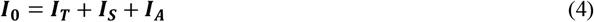

Where ***I*_0_**, ***I_T_, I_S_*** and ***I_A_*** are the incident light intensity, transmittance, scattering (including reflection) and absorbance intensity, respectively. For a blank solvent, e.g. water, *I_S_* is assumed to be zero, and the above equation can be rewritten as Eq. 5.

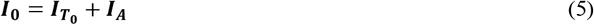

Where ***I*_*T*_0__** is the transmittance of water and then it can be rewritten as Eq. 6.

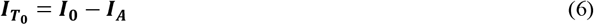

For particle suspension, the scattering intensity can be expressed as Eq. 7 by combining both Eq. 4 and Eq. 6.

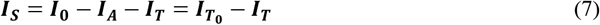

According to Beer-Lambert law, 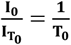 while 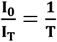, and the measured scattering intensity (*I_S,m_*) in Eq. 7 can be expressed as

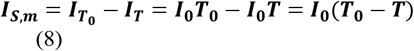

where ***T*_0_** and *T* is the transmittance of the solvent before and after the particle dispersion, respectively. Assuming the theoretical scattering intensity is equal to the measured one, i.e. *I_S,t_*=*I_S,m_*, a new relationship can be obtained combining Eq. 3 and Eq. 8.

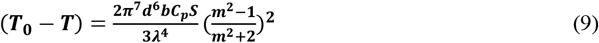

Through the use of reference materials (RM) with certain size (*d_s_*), concentration (*C_p_*), refractive index (*m*) and the known optical path length (*b*), Eq. 9 can be further simplified as

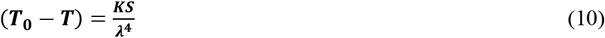

where 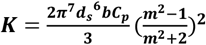, is a constant for a given monodispersed particle suspension. Therefore, the illumination spot area, *S* can be determined by fitting 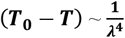 in Eq. 10. Finally, the above equation can be rewritten as

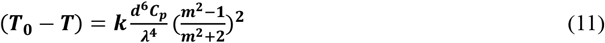

as long as *S* is determined, where 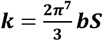. Thus, the diameter (*d*) of the particles in a suspension can be measured by the change of their transmittance at any wavelength according to Eq. 11 if *C_p_* and *m* are known.

### Material

The polystyrene (PS) nanoparticles and the Bovine serum albumin (BSA) used in this work were purchased from Sigma-Aldrich The polystyrene nanoparticles were amine-modified and labelled as L0780-1ML with a nominal diameter of 50 nm (named PS50) and L9904-1ML of 100 nm (named PS100), respectively, and their average density is 1.05 g/cm^3^ and solid content is 2.5 wt%. The polystyrene RM with a verified diameter of 79.1±1.5 nm was manufactured by the China University of Petroleum. The silica nanoparticles were synthesized by a modified Stöber method. The polystyrene spheres and the silica nanoparticles were diluted with the ultrapure water to different concentration prior to use, and then divided into two parts, the one was used as the reference and another one is incubated with BSA for 12 hours prior to measurement. These samples are labelled as PS50, PS50@BSA, PS100, PS100@BSA, SiO_2_, SiO_2_@BSA, respectively.

The UV-Vis transmission spectra were measured in a quartz glass cuvette with a path length of *b*=1 cm, and filled with deionized water as the blank sample by a miniaturized spectrometer (Perkin Elmer, Lambda 650). The spectra were recorded in wavelength range of 400<λ<600nm and the wavelength interval is set to 1 nm. All measurements were carried out at room temperature. For comparison, FEI Tecnai F20(Hillsboro, America), Anton Paar Litesizer 500 Dynamic Light Scattering (Graz, Austria)and Carle Zeiss Merlin high resolution thermal field emission scanning electron microscope(Oberkochen, Germany) was used to measure the hydraulic and ‘dry’ diameter of the polystyrene and silica nanoparticles. And the attenuated total reflectance (ATR) infrared spectroscopy(Thermo Fisher Scientific, Waltham, MA) was used to analyse the adsorbed protein conformation.

## RESULTS AND DISCUSSION

Figure 1 shows the image of SiO_2_ NPs and their BSA shell by the cryogenic transmission electron microscope (Cryo-TEM), it can be seen that the protein has been adsorbed on the surface of the nanoparticles and formed a uniform adlayer around the particles. The average thickness of the protein shell on the nanoparticles is ca. 1.5 nm. Figure 2 shows the SEM images of the PS and silica nanoparticles before and after the protein adsorption with uniform size. When the BSA is absorbed on the nanoparticles, the observed images are more charging due to the formation of the less conductive protein adlayer on these particles. According to the statistical results based on SEM images, the size of PS50, PS50@BSA, PS100, PS100@BSA, SiO_2_, SiO_2_@BSA are 45.85, 50.42, 74.59, 75.85, 38.80, and 42.64 nm, respectively. Therefore, the thickness of the protein adlayer is determined as 4.57, 1.26 and 3.84 nm for PS50@BSA, PS100@BSA and SiO_2_@BSA, respectively.

**Figure 1.**
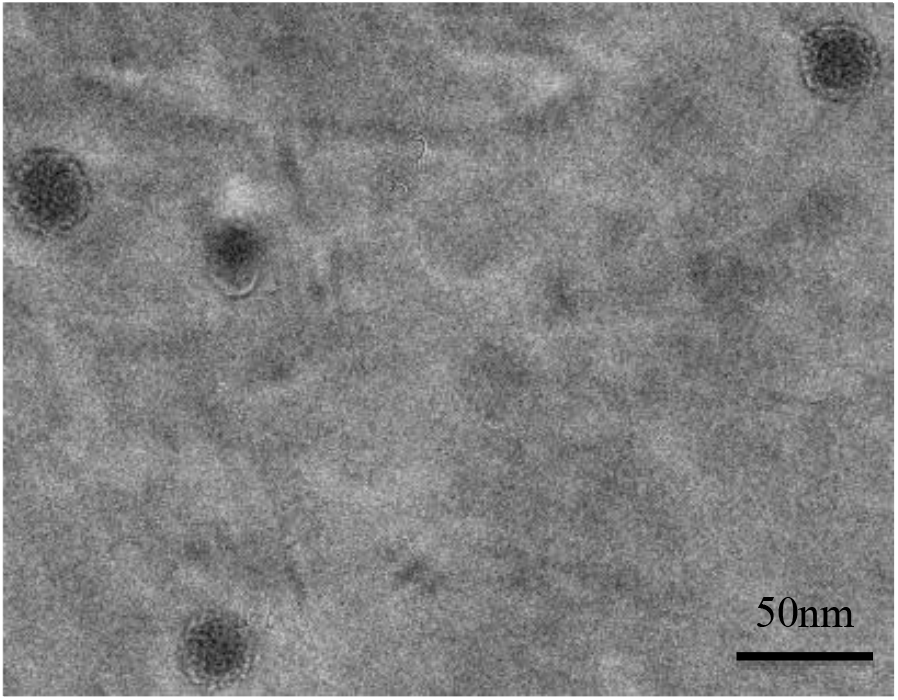
Cryo-transmission electron microscope image of SiO_2_@BSA

**Figure 2.**
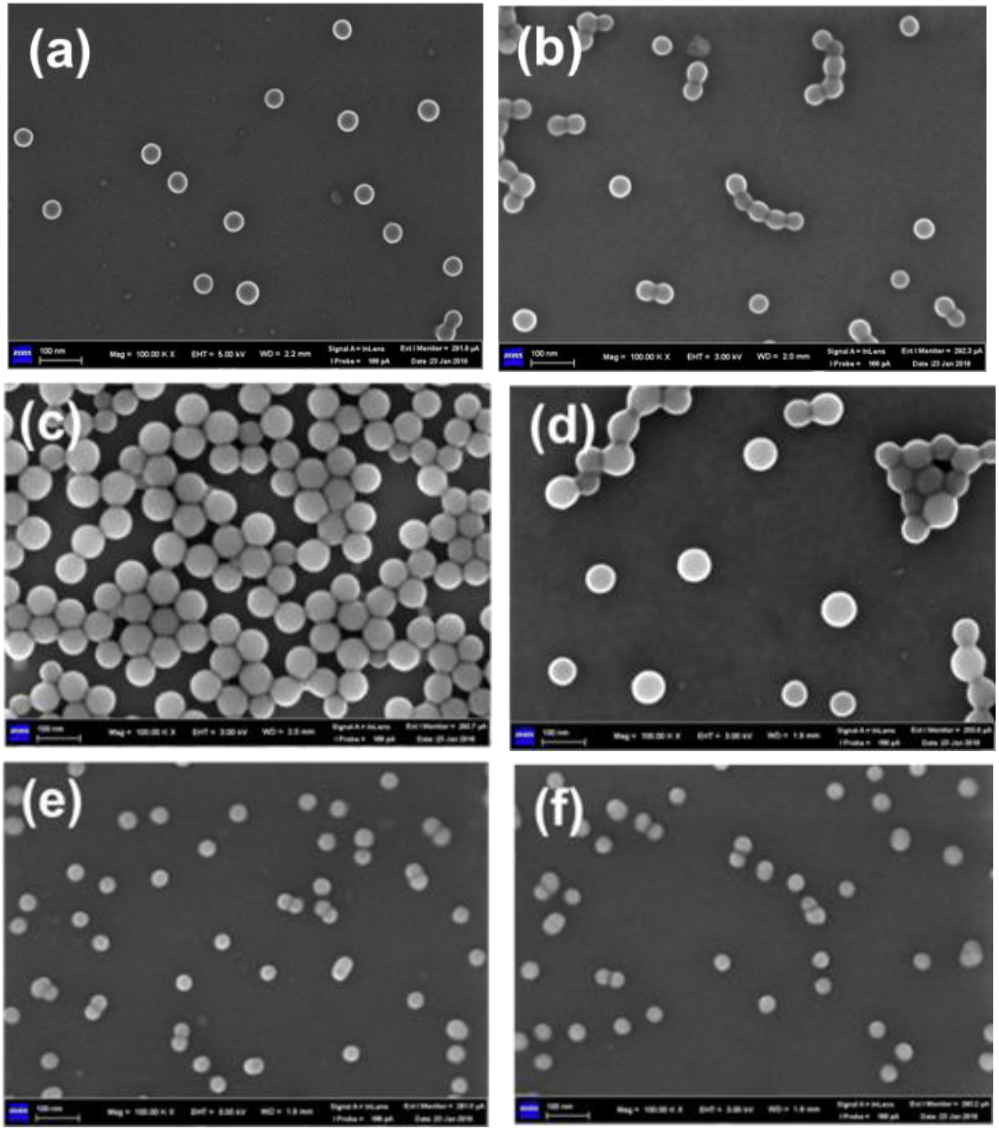
Scanning electron microscopy images of a) PS50, b) PS50B, c) PS100, d) PS100B, e) SiO_2_ and f) SiO_2_@BSA

Table S1 shows the dynamic light scattering data of the nanoparticles (see the Supporting Information). The curves show that the size distribution of the nanoparticles conforms to Log-Normal distribution and the particles are monodispersed. As it can be seen that the samples absorbed with BSA are obviously larger than their corresponding bare particles. According to the Z-average size results measured by dynamic light scattering, the size of the samples is 54.38, 58.24, 121.54, 124.21, 71.70, 75.73 nm, for PS50, PS50@BSA, PS100, PS100@BSA, SiO_2_ and SiO_2_@BSA, respectively. The thickness of the protein adsorption layer is 3.86 nm, 2.67 nm and 4.03 nm for PS50@BSA, PS100@BSA, SiO_2_@BSA.

Both the PS microspheres and BSA have no characteristic absorption in the visible range of 400-600 nm, as shown in Figure S1 (see Supporting information). Then the transmittance loss is considered to be caused by the light scattering by particles. The diameter of the particles can be calculated according to Eq. 11 By RGDA. However, in order to determine the size of the particle with unknown diameter, the spot parameter S must be determined in advance assisted by the use of the PS reference particles (79.1±1.5 nm). Figure 3 validates the applicability of the RGDA method, where the diameter of the reference particles determined by RGDA is almost constant with the irradiation wavelength.

**Figure 3.**
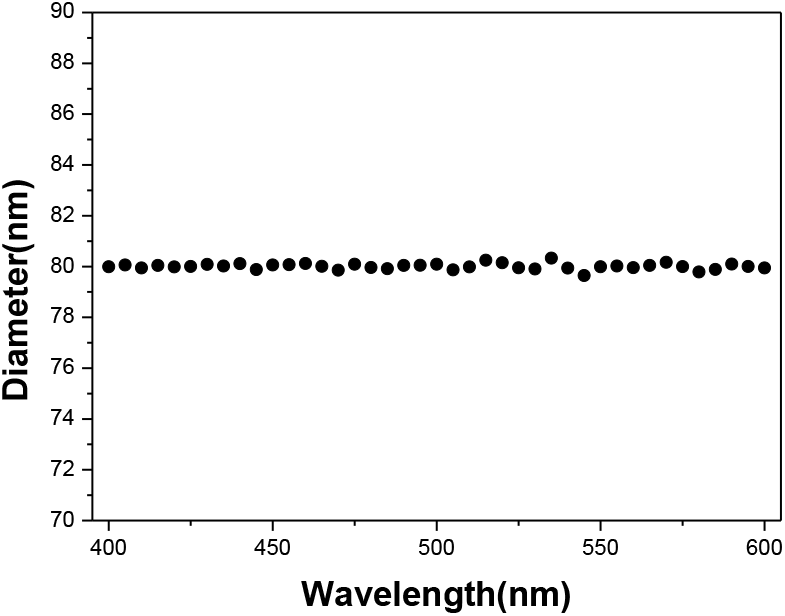
The diameter of the reference PS particles of 80 nm calculated by RGDA

The polystyrene and silica particle size calculated by RGDA is shown in Figure 4-6. The diameter of the samples calculated by RGDA method decreases gradually with the increase of the irradiation wavelength. We tentatively attribute this reason to the wavelength-dependent refraction. For a particle with given size, the longer the wavelength, the stronger the refraction intensity is. It, therefore, causes the underestimate on the scattering intensity and the particle size determined by the above equations, particularly for the smaller particles, PS50 and SiO_2_ as shown in Figure 4 and 6. As can be seen that the calculated diameter of the PS100 and PS100@BSA has a big lump occurs centred at 490 nm (Figure S2, Supporting Information), which can be contributed to the additional adsorption caused by the fluorescent dye encapsulated in the commercial product PS100. The product specification shows that these commercial particles can be exited at around 520 nm (absorbance). Therefore, we apply the baseline correction in order to eliminate the additional absorption caused by the fluorescent molecules in PS100, and the corrected result is shown in Figure 5. The data points for silica nanoparticles become more discretized as seen in Figure 6 compared with the other samples due to bad signal-to-noise ratio. This could be attributed to the lower relative refractive index (m) of silica NPs compared with that of PS materials, and although the silica particle concentration is higher, but the size is smaller than the PS materials. As can be seen that the diameter determined by RGDA changes little with the different wavelength. However, the size calculated by RGDA is much closer to those determined by SEM or TEM than those measured by DLS and the best-matched results are obtained at 400 nm, indicating that the diameter measured by RGDA is a ‘dry’ size not a hydraulic one as DLS does.

**Figure 4.**
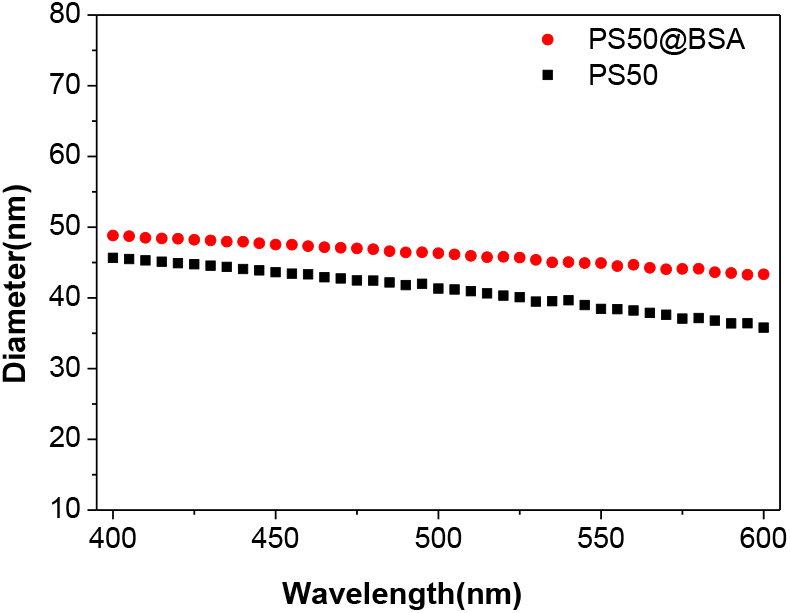
The diameter of PS50 and PS50@BSA calculated by RGDA

**Figure 5.**
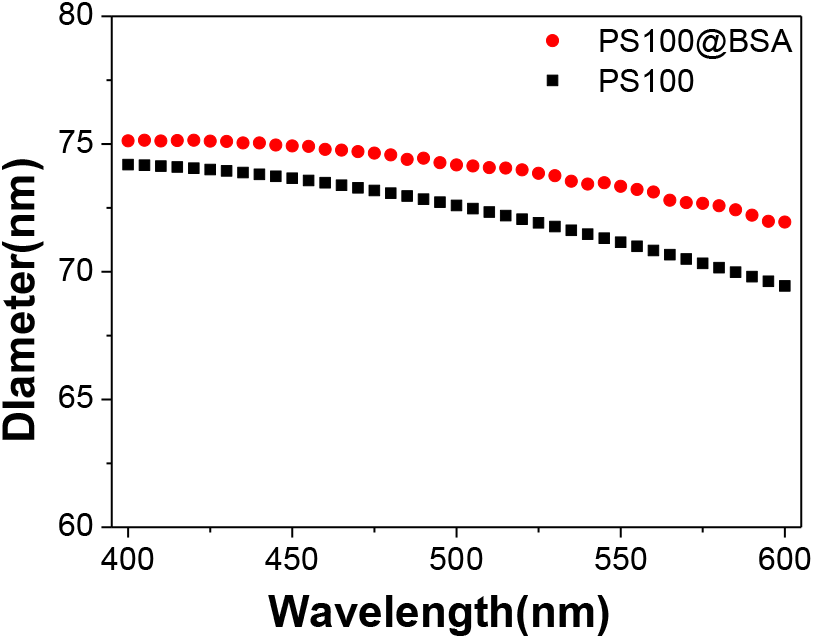
The diameter of PS100 and PS100@BSA calculated by RGDA

**Figure 6.**
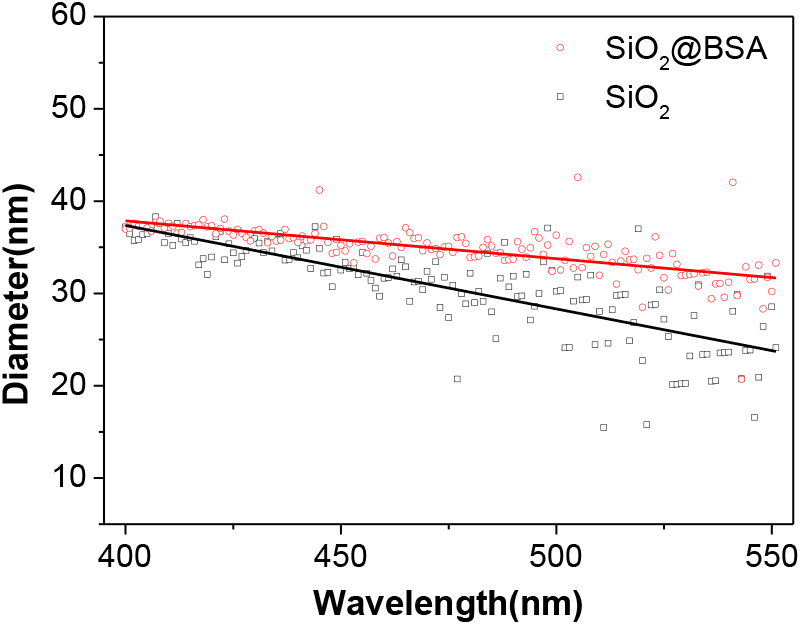
The diameter of the SiO_2_ and SiO_2_@BSA calculated by RGDA

Figure 7 shows the calculated adlayer thickness for the BSA molecules on PS50, PS100 and SiO_2_ particles increased with the increase of the irradiation wavelength. It shows that the adlayer adsorbed on PS50 is thicker than that on PS100 and SiO_2_ NPs. This might be attributed to the conformation change of the protein molecules absorbed on the particles with different curvature. Perry et al have studied the adsorption of the BSA and fibrinogen onto both hydrophilic and hydrophobic substrates of varying sizes, and found that the saturated adsorption amount for BSA onto nanoparticles maximizes at ~40 nm accompanying the change of their secondary structure.(17) The albumin absorbed on the larger nanoparticles loses more helix structure than the smaller particles and less closely-packed adlayer is formed, which corresponds to the formation of thinner BSA layer on PS100 than PS50.

**Figure 7.**
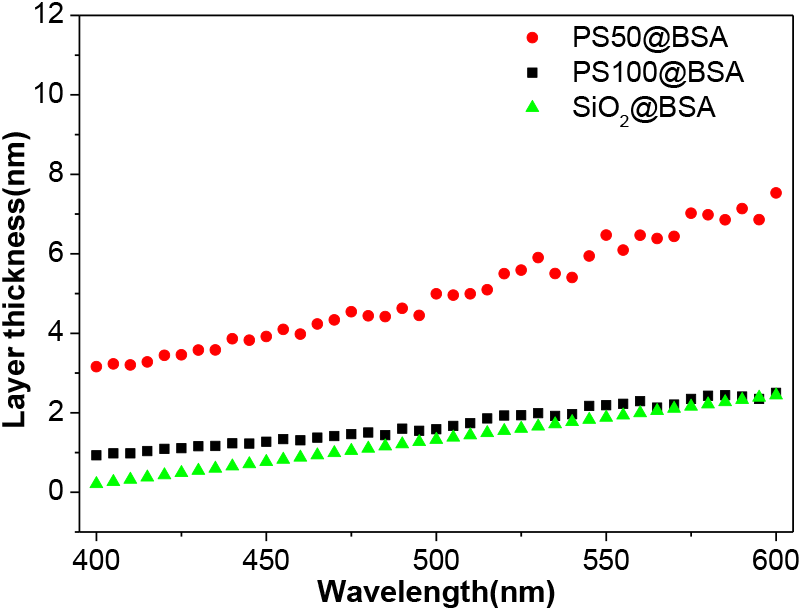
The layer thickness of the PS50@BSA, PS100@BSA and SiO_2_@BSA calculated by RGDA

Figure 8 shows the BSA, PS50@BSA, PS100@BSA infrared spectra mainly from amide bond vibrations. The amide I band centred at 1700-1600 cm^−1^ is largely due to C=O stretching vibrations. This band is sensitive to the change in secondary structure and has therefore been widely used for protein conformational analysis. As seen in Figure 8, the peaks at 1682 and 1667 cm^−1^ have been assigned to β-turn structures, 1658 and 1649 cm^−1^ to α-helixes, 1642 cm^−1^ to random chains, 1634 and 1619 cm^−1^ to extended chains or β-sheet, and ~1610 cm^−1^ are considered to arise from intermolecular bonding. Compared the FT-IR spectra, it can be seen that the disappearance of many amide I peaks, particularly, the peak at 1649 cm^−1^ in PS100@BSA indicates that the loss of its secondary structure, especially α-helix, The corresponding peaks are preserved in PS50 and BSA, demonstrating that the BSA molecules adsorbed on smaller particles or in their free state can largely keep their secondary structure and those absorbed on larger particles undergo severe structure deformation. The larger structure deformation or loss in PS100 makes the adlayer thickness smaller while the less deformed BSA shell on PS50 largely possesses its original dimension as it in free state, where the three dimensions of the BSA molecule are determined as 14.1±0.5, 4.2±0.4 and 4.2±0.4, respectively.(18) The size of BSA is quite close to the thickness of the adlayer on PS50 determined by both RGDA and SEM as seen in Table 1. However, due to the existence of a great number of Si-OH groups on SiO_2_ NPs, the interaction of albumin-SiO_2_ is considered stronger than that of albumin-PS of same size and, therefore, the heavier deformation causes thinner adlayer on SiO_2_ NPs as seen in Figure 7 and Table 1.

**Figure 8.**
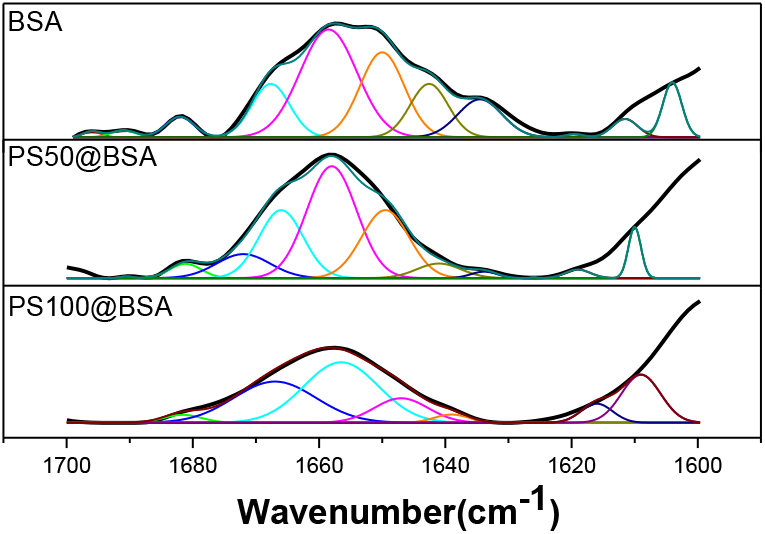
FT-IR spectra for BSA, PS50@BSA, PS100@BSA and the fitted component peaks

**Table 1.** Colloidal nanoparticles size and their BSA adlayer thickness

| method | size | PS50 |  | PS100 |  | SiO<sub>2</sub> |  |
| --- | --- | --- | --- | --- | --- | --- | --- |
|  |  | bare | @BSA | bare | @BSA | bare | @BSA |
| **RGDA**<br>(nm) | NP Diameter | 45.5 | 48.8 | 74.2 | 75.1 | 36.9 | 38.0 |
|  | Adlayer thickness |  | 3.3 |  | 0.9 |  | 1.2 |
| **SEM**<br>(nm) | NP Diameter | 45.8 | 50.4 | 74.6 | 75.8 | 38.8 | 40.1 |
|  | Adlayer thickness |  | 4.6 |  | 1.3 |  | 1.3 |
| **DLS**<br>(nm) | NP Diameter | 54.4 | 58.2 | 121.5 | 124.2 | 71.7 | 75.3 |
|  | Adlayer thickness |  | 3.9 |  | 2.7 |  | 4.0 |

Table 1 summarize and compare the size and dimension results measured by RGDA, DLS and SEM, respectively. It can be seen clearly that the diameter determined by RGDA is quite close to that measured by electron microscopy while significant smaller than that measured by DLS since the size determined by DLS contains the a few nanometers of the hydrated layer around the particles. The size determined by RGDA in short wavelength is closer to that observed by SEM, e.g. 400 nm. Therefore, all the RGDA values listed in Table 1 are calculated using 400 nm as its irradiation wavelength.

Compared with conventional DLS, the RM assisted RGDA is able to measure the colloidal particles in a more sensitive (e.g. very diluted solution), economic and convenient way, where, as a function of *λ*, the determination of the effective, wavelength-dependent and instrument-related irradiation spot area is a key. The spot area S has been considered as a constant in previous researches while we found that it is a function of *λ* in this study, which can be accurately determined by the use of reference materials with verified size and specific refractive index. However, one of the highlighted features for RGDA is its ability to measure the ‘dry’ particle size in situ in solution not like DLS, which is inevitably disturbed by the hydrated layer on the particle surface. In addition, the thickness of the protein adlayer adsorbed on either polymer or inorganic NPs can be directly measured by this method. Finally, this method is applicable in a very wide range of wavelength not like DLS, which can only work at a fixed wavelength, e.g. 658 nm for many DLS machines, and avoid the disturbance and overlapping of the light spectra from the unexpected absorbance and emission at specific wavelength, e.g. some fluorescent quantum dots or noble metal nanocrystals with significant surface plasmon resonance which are difficult to be measured by light scattering method. Further studies on this issue will be conducted in near future.

## CONCLUSIONS

Reference materials assisted RGDA (rm-RGDA) has been developed and used to measure the size and thickness of the adlayer on the particles in situ in solution. The particle size determined by rm-RGDA is quite close to that measured by electron microscopy but significantly smaller than that measured by DLS. The BSA adlayer absorbed on PS50, PS100 and SiO_2_ NPs is 3.3, 0.9 and 1.2 nm, respectively, and close to those observed by SEM, which is 4.6, 1.3 and 3.8 nm, respectively. The FTIR analysis results show that the BSA absorbed on flat, e.g. larger PS100 particles or hydroxyl-abundant surface, e.g. SiO_2_ NPs can lose its secondary structure, e.g. α-helix, to a great extent and that absorbed on a more curve surface, e.g. smaller PS50 particles can largely preserve its secondary structure like it in free state. These dimension measurement results are closely related to their structure change and those reported in previous literatures. We believe this method provide a facile and sensitive way to measure the dimension and its absorption-dependent change of the hybrid and core-shell structured nanoparticles in a wide range of wavelength.

## AUTHOR CONTRIBUTIONS

Performed the experiments: LY; conceived the project: GG; planed the work: CL; developed the model: LY, CL; analysed the data: DL, XG, ZZ; wrote the manuscript: ZZ, LY

## ACKNOWLEDGMENTS

The authors thank the National Key Research and Development Program of China, (Grant No. 2016YFA0200904) for financial support.

## REFERENCES

1 Apfel, U.; Horner, K. D.; Ballauff, M., Precise Analysis of the Turbidity Spectra of a Concentrated Latex. Langmuir 1995, 11(9), 3401–3407.

2 Crawley, G.; Cournil, M.; DiBenedetto, D., Size analysis of fine particle suspensions by spectral turbidimetry: Potential and limits. Powder Technol 1997, 91(3), 197–208.

3 Shmakov, S. L., Allowance for instrumental factors in the turbidity and light-scattering spectrum methods. Optics and Spectroscopy 2003, 95(3), 461–463.

4 Haiss, W.; Thanh, N. T. K.; Aveyard, J.; Fernig, D. G., Determination of size and concentration of gold nanoparticles from UV-Vis spectra. Analytical Chemistry 2007, 79(11), 4215–4221.

5 Khlebtsov, B. N.; Khanadeev, V. A.; Khlebtsov, N. G., Determination of the size, concentration, and refractive index of silica nanoparticles from turbidity spectra. Langmuir 2008, 24(16), 8964–8970.

6 Pramanik, S.; Banerjee, P.; Sarkar, A.; Bhattacharya, S. C., Size-dependent interaction of gold nanoparticles with transport protein: A spectroscopic study. Journal of Luminescence 2008, 128(12), 1969–1974.

7 Lechner, M. D.; Colfen, H.; Mittal, V.; Volkel, A.; Wohlleben, W., Sedimentation measurements with the analytical ultracentrifuge with absorption optics: influence of Mie scattering and absorption of the particles. Colloid and Polymer Science 2011, 289(10), 1145–1155.

8 Glasse, B.; Assenhaimer, C.; Guardani, R.; Fritsching, U., Turbidimetry for the Stability Evaluation of Emulsions Used in Machining Industry. Canadian Journal of Chemical Engineering 2014, 92(2), 324–329.

9 Wang, Y. M.; Steinhoff, B.; Brinkmann, C.; Alig, I., In-line monitoring of the thermal degradation of poly(L-lactic acid) during melt extrusion by UV-vis spectroscopy. Polymer 2008, 49(5), 1257–1265.

10 Khlebtsov, B. N.; Kovler, L. A.; Bogatyrev, V. A.; Khlebtsov, N. G.; Shchyogolev, S. Y., Studies of phosphatidylcholine vesicles by spectroturbidimetric and dynamic light scattering methods. J Quant Spectrosc Ra 2003, 79, 825–838.

11 Greengard, L.; Hagstrom, T.; Jiang, S. D., Extension of the Lorenz-Mie-Debye method for electromagnetic scattering to the time-domain. J Comput Phys 2015, 299, 98–105.

12 Elsayed, M. M. A.; Cevc, G., Turbidity Spectroscopy for Characterization of Submicroscopic Drug Carriers, Such as Nanoparticles and Lipid Vesicles: Size Determination. Pharm Res-Dordr 2011, 28(9), 2204–2222.

13 Guschin, V.; Becker, W.; Eisenreich, N.; Bendfeld, A., Determination of the Nanoparticle Size Distribution in Media by Turbidimetric Measurements. Chem Eng Technol 2012, 35(2), 317–322.

14 Glasse, B.; Riefler, N.; Fritsching, U., Intercomparison of Numerical Inversion Algorithms for Particle Size Determination of Polystyrene Suspensions Using Spectral Turbidimetry. J Spectrosc 2015, 1–10.

15 Le Blevennec, G., Detection limits for nanoparticles in solution with classical turbidity spectra. Instrumentation, Metrology, and Standards for Nanomanufacturing, Optics, and Semiconductors Vii 2013, 8819.

16 Zhao, H. Y.; Brown, P. H.; Schuckt, P., On the Distribution of Protein Refractive Index Increments. Biophys J 2011, 100(9), 2309–2317.

17 Roach, P.; Farrar, D.; Perry, C. C., Surface tailoring for controlled protein adsorption: Effect of topography at the nanometer scale and chemistry. J Am Chem Soc 2006, 128(12), 3939–3945.

18 Wright, A. K.; Thompson, M. R., Hydrodynamic Structure of Bovine Serum-Albumin Determined by Transient Electric Birefringence. Biophys J 1975, 15(2), 137–141.

